# Culture wars: Empirically determining the best approach for plasmid library amplification

**DOI:** 10.1101/2024.05.24.595835

**Authors:** Nicholas Mateyko, Carl de Boer

## Abstract

DNA libraries are critical components of many biological assays. These libraries are often kept in plasmids that are amplified in *E. coli* to generate sufficient material for an experiment. Library uniformity is critical for ensuring that every element in the library is tested similarly, and is thought to be influenced by the culture approach used during library amplification. We tested five commonly used culturing methods for their ability to uniformly amplify plasmid libraries: liquid, semisolid agar, cell spreader-spread plates with high or low colony density, and bead-spread plates. Each approach was evaluated with two library types: a random 80-mer library, representing high complexity low coverage of similar sequence lengths, and a human TF ORF library, representing low complexity high coverage of diverse sequence lengths. We found that no method was better than liquid culture, which produced relatively uniform libraries regardless of library type. However, when libraries were transformed with high coverage, culturing method had minimal impact on uniformity or amplification bias. Plating libraries was the worst approach by almost every measure for both library types, and, counter-intuitively, produced the strongest biases against long sequence representation. Semisolid agar amplified most elements of the library uniformly but also included outliers with orders of magnitude higher abundance. For amplifying DNA libraries, liquid culture, the simplest method, appears to be best.

## Introduction

DNA libraries are a crucial component of a wide range of biological experiments, including cDNA library screening^1^, CRISPR screens^2^, massively parallel reporter assays^3^, deep mutational scans^4^, and directed evolution of proteins^5^. These libraries are usually constructed in plasmids that are transformed into a microorganism such as *Escherichia coli* (*E. coli*); these transformants are then grown to amplify the plasmid library, and the amplified DNA extracted to complete the experiment^6^. As the transformants replicate, different clones may grow at different rates, changing the relative abundance of the sequences present in the library. This uneven amplification is undesirable for many applications, as it results in the inability to characterize lowly abundant sequences, which may even be altogether absent from the final experiment (“dropping out”), while highly abundant sequences waste a large number of sequencing reads or use up screening resources that would be better spent on other sequences. Many new methods require extremely complex plasmid libraries^7,8^, and the size of libraries will likely continue to grow with increased sequencing throughput. Given the investment required to generate and screen large plasmid libraries, it is crucial to ensure that each unique sequence remains at the desired abundance after library amplification so that as many high-quality measurements as possible can be made.

Liquid culture is the simplest method for amplifying plasmids in *E. coli* and is the standard technique for amplifying a single plasmid species. However, many publications assert that amplifying plasmid libraries in liquid culture can result in very non-uniform amplification, as a small number of fit clones can take over the culture while other clones end up at such low abundance that they are lost upon sampling of the library^6,9,10^. To address this issue, many protocols call for amplifying plasmid libraries on agar plates, which supposedly reduces competition between clones by limiting the local nutrients available to each colony^11,12^. However, it is difficult to evenly spread colonies on a plate, and the number of plates required becomes unreasonable for very high complexity libraries. Other protocols use a semisolid agarose medium that combines the benefits of both liquid and solid amplification: colonies are spread evenly throughout the medium yet are still limited by their local nutrient supply, and laborious spreading and colony scraping are not required^6,9,10^. Despite the criticality of the library culturing approaches to the experimental outcome, to our knowledge, no direct comparisons of their abilities to uniformly amplify libraries has yet been performed.

Here, we sought to test the efficacy of liquid, solid, and semisolid culture methods to uniformly amplify plasmid libraries. To reliably quantify bias resulting from culture methods, we use two paradigms where we can reliably quantify the input library complexity, either because the starting complexity is so high that there can be at most one of each library element, or because the library already exists and we are simply re-transforming it (**Fig. 1**). The two library types we use are representative of those commonly used in modern research methods: transgenes of diverse lengths that may impose metabolic or other fitness costs (human transcription factors), and short (80 bp) random sequences that are unlikely to affect fitness substantially. We find that liquid culture, the simplest method, performs at least as well as other methods when complexity is very high and transformation coverage is low (1X coverage). However, when library complexity is low and transformation coverage is high (100X coverage), culture methods have minimal impact on library uniformity and do not detectably bias library composition by element length. Our results will enable researchers to make evidence-based decisions for how they amplify their plasmid libraries so they can generate the highest quality data in their experiments without wasting time and resources.

**Figure 1.**
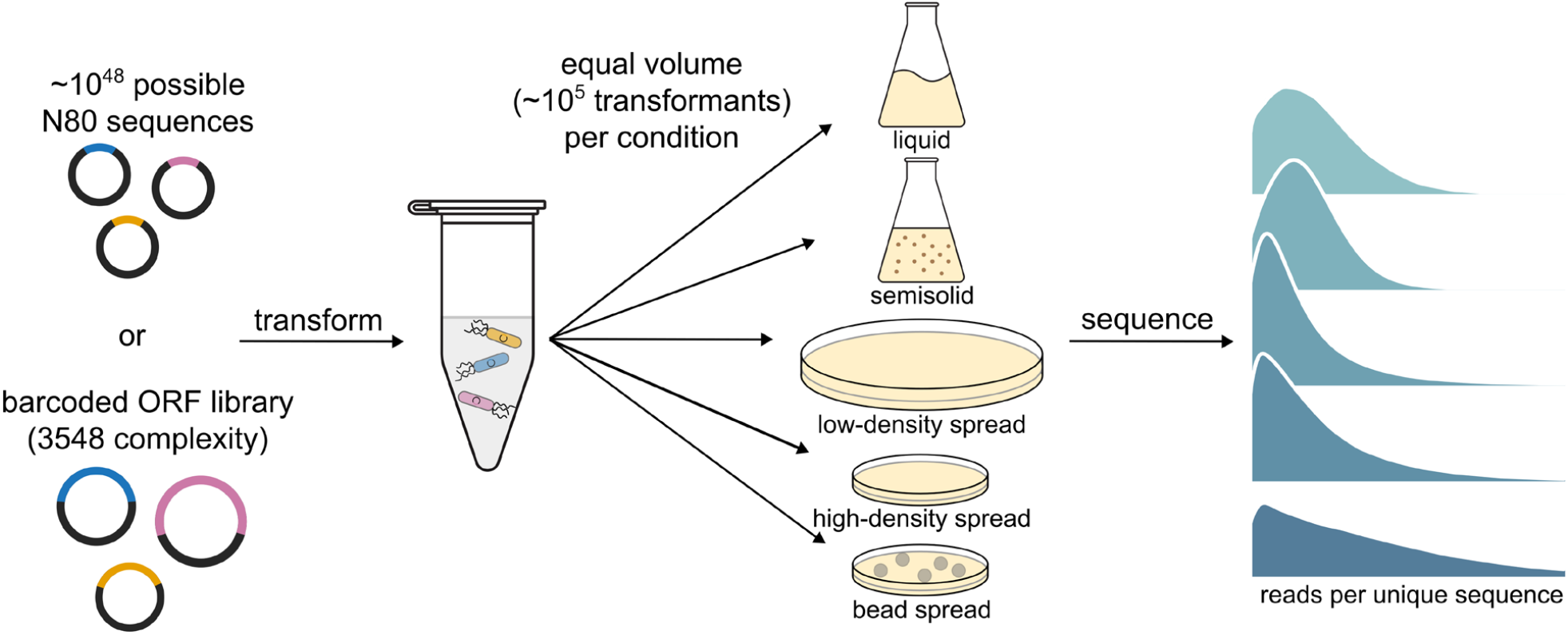
Experimental procedure. A plasmid library containing only a single copy of each unique random sequence or a barcoded library of human transcription factor ORFs was transformed into *E. coli*. Equal volumes of recovered transformants were grown in five different culture conditions. Each plasmid library was sequenced to assess how evenly the library was amplified.

## Results

### Liquid culture amplifies plasmid libraries most uniformly

We first wanted to test the scenario of a highly complex library with low transformation coverage, where random differences between clone growth could lead to noisy amplification of the library and large differences in the proportion of library members. We chose to use a library in which each sequence was at an equal initial abundance of 1, such that each sequence could be used as a barcode for a single transformant clone. To accomplish this, we cloned a fragment of DNA containing 80 random bases (N80) into a plasmid and transformed it into *E. coli*. Because the random inserts were never amplified before transformation and the number of potential starting molecules is so much higher than the number cloned (4^80^ ≈ 10^48^), it is highly improbable that any sequence was present more than once at transformation. This library was then transformed into *E. coli* and each transformation was divided equally and grown using different culture methods (**Fig. 1)**. For the N80 experiment, we aimed for ∼100,000 transformants for each separately cultured library.

We tested several commonly used culturing techniques. Three grew the library as colonies on top of a solid agar medium, which is supposed to limit competition between different clones as each colony restricts its own growth. Using a standard glass cell spreader, we plated transformants at a high density of colonies (∼1200/cm^2^) to simulate a density that would be reasonable for libraries on the order of 10^7^ complexity, as well as at ∼1/8 this density to see if increased space between colonies reduces competition. We also tested high colony density spread with glass beads, which tend to produce more evenly spread colonies than cell spreaders. Finally, we tested semi-solid agarose, which gels after being mixed with the transformed cells, ensuring that colonies grow suspended evenly throughout the entire volume. We compared each culturing approach to liquid culture, which, being the easiest approach, is the standard to beat.

Next, we compared library element abundance by high-throughput sequencing. After extracting and sequencing the plasmid libraries, we subsampled reads to equal depth, counted the number of unique sequences in each library and the number of reads per unique sequence, then compared each culture method to liquid culture. As equal volumes were used for each culture method derived from the same recovered transformation, all libraries were expected to contain a similar number of unique sequences. Surprisingly, there was significant variability in the number of observed sequences between culture methods, with liquid culture having the fewest, plate-based methods having about 7% more than liquid culture, and semisolid having about 35% more than liquid culture (**Fig. 2a**).

**Figure 2.**
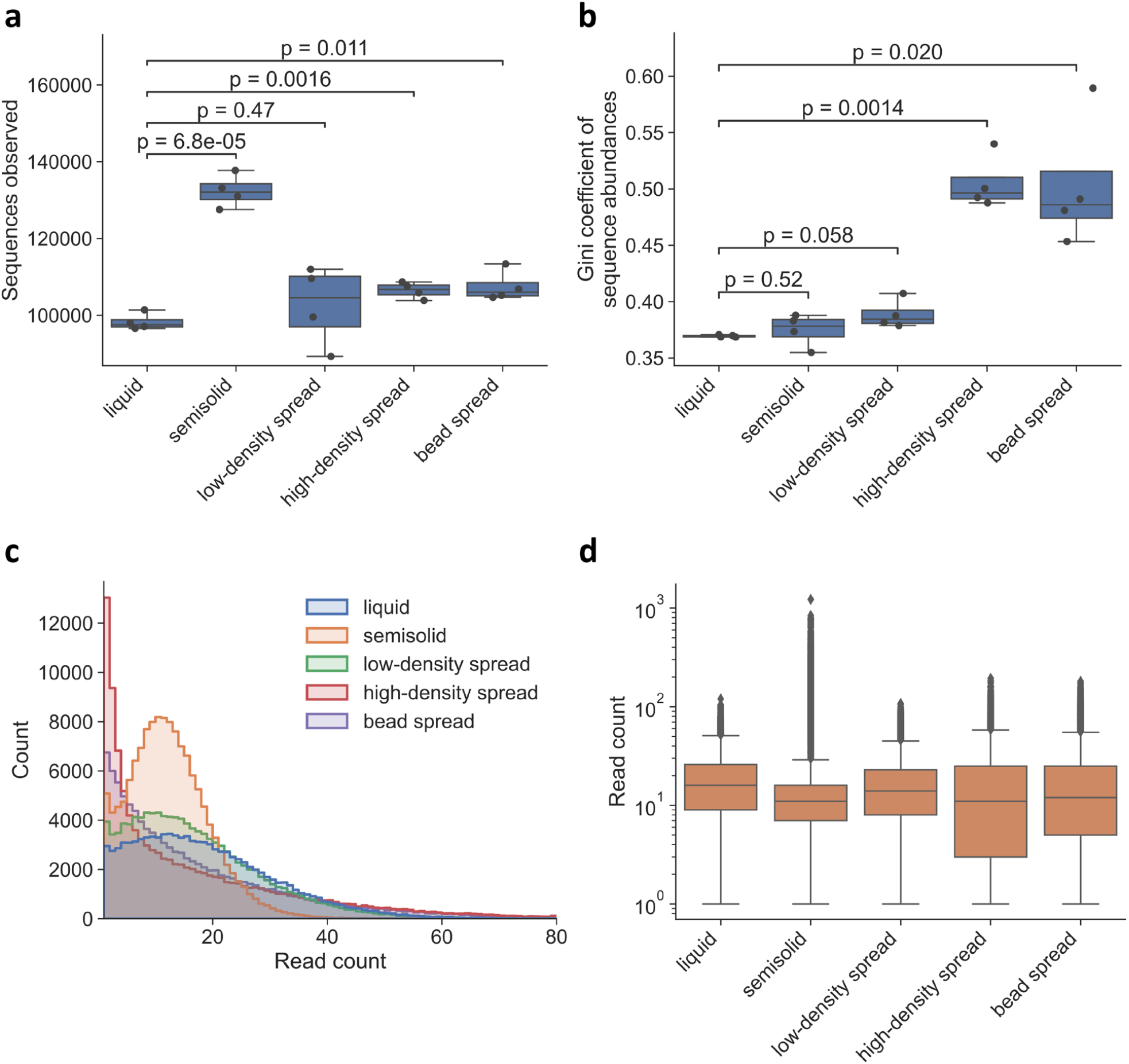
Liquid culture yields the most uniform but lowest complexity libraries. **a**. Comparison of the number of unique N80 sequences observed in each library (*y* axis) for each culture method (*x* axis). **b**. Comparison of library uniformity, measured by the Gini coefficient of sequence abundances (*y* axis) for each culture method (*x* axis). **c**. Representative replicate showing the distribution of sequence counts (*y* axis) with the corresponding number of reads (*x* axis) for each culture condition (colours). **d**. Distributions of read counts per sequence (*y* axis; log scale) for each culture condition (*x* axis), showing outliers in the semisolid condition (representative replicate). Box plot boxes represent the median and interquartile range (IQR). Whiskers extend to points within 1.5 IQRs of the first and third quartiles. P-values were calculated using Welch’s t-test.

We then compared the uniformity of amplification for each library amplification method to liquid culture. Each library started out with an equal abundance of each sequence, so a perfectly uniformly amplified library would retain this equal abundance. A good measure of this uniformity is the Gini coefficient, which ranges from 0 to 1, where a library with equal proportions of all sequences would have a coefficient of 0. No culture method was better than liquid culture by this measure (**Fig. 2b**). High-density spread and bead spread methods had significantly higher Gini coefficients than liquid culture, indicating less uniform amplification of the library, while the other methods were not significantly different than liquid culture (**Fig. 2b**). Although the semisolid read-count distribution appeared less disperse than others (**Fig. 2c**), it contained many outlier sequences with much higher read counts than seen in the other methods (**Fig. 2d**).

### High transformation coverage mitigates culturing bias in libraries of diverse length

We next sought to test the culture methods when amplifying a library that is representative of heterogeneous cDNA or ORF libraries in which the length of library elements can span several orders of magnitude. Because the library is much more heterogeneous, there are likely to be more substantial effects on fitness between clones, which could manifest as increasing bias when culturing the library. We chose the Multiplexed Overexpression of Regulatory Factors (MORF) library^13^ for this purpose, which contains 3548 human transcription factor isoforms which range in length from 150 bp to 9690 bp and includes a barcode sequence enabling quantification of each library element by targeted sequencing of the barcode region. This library was transformed into *E. coli* and grown with the same set of culture conditions as the N80 library, with ∼100X library coverage for each culture. Plasmids were extracted and barcodes from each cultured library and the input plasmid library were amplified and sequenced so that the change in library composition could be measured for each culture method.

Unlike the previous case of a library with low transformation coverage, there were few robust differences between the culture methods (**Fig. 3a-c**). As expected given the high transformation coverage, most library elements were seen in each replicate and the number of elements observed was similar regardless of culturing method (**Fig. 3a**). Again, no culturing method had a better uniformity than liquid culture (**Fig. 3b,c**) and all of them appeared to maintain the original composition equally well (**Fig. 3d)**.

**Figure 3.**
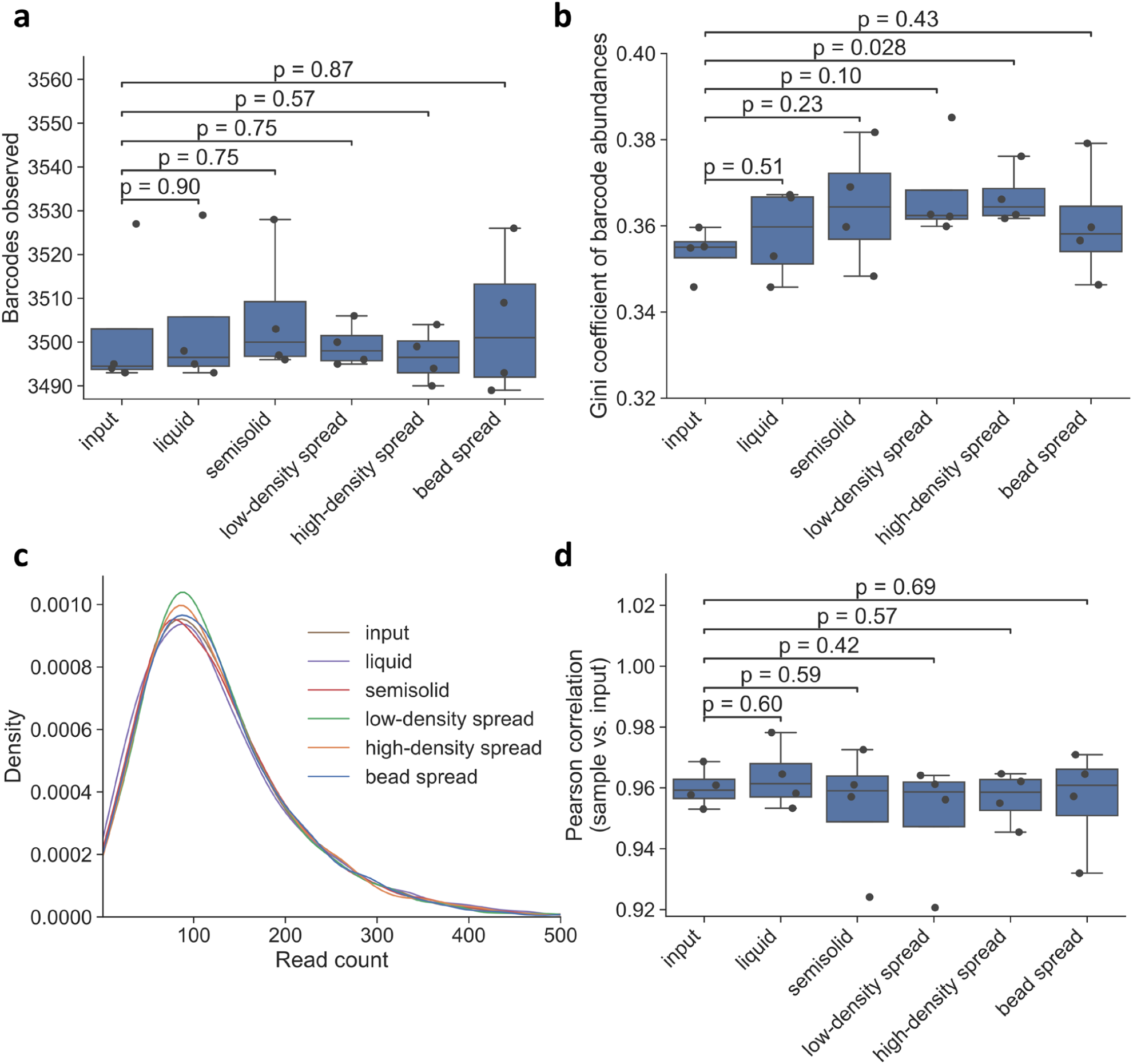
Distributions and statistics of read counts for MORF libraries. **a**. Comparison of the number of unique ORF barcodes observed in each library (*y* axis) for each culture method (*x* axis). **b**. Comparison of library uniformity, measured by the Gini coefficient of barcode abundances (*y* axis) for each culture method (*x* axis). **c**. Representative replicate showing the distribution of sequence counts (*y* axis) with the corresponding number of reads (*x* axis) for each culture condition (colours). **d**. Pearson correlation between read counts for each sample and input read counts (*y* axis) for each culture condition and the input library (*x* axis). For each replicate, the Pearson correlation between barcode counts for each sample and the average input barcode counts for the other three replicates was calculated. Box plot boxes represent the median and interquartile range (IQR). Whiskers extend to points within 1.5 IQRs of the first and third quartiles. P-values were calculated using Welch’s t-test.

We then looked at whether certain culture methods specifically biased against amplification of larger library elements. Since larger plasmids could potentially be a larger burden on their host cells, for instance due to their metabolic cost or biochemical activities encoded within them, the growth rates between library elements could differ. If this significantly biased library composition, we would expect to see the most substantial effect for liquid culture, where all clones are in direct competition for the same pool of nutrients. The MORF library already had a substantial bias towards shorter insert sizes in the input library (**Fig. 4a**), potentially due to the Gateway cloning step used to create it. However, the length bias was not significantly changed by any of the culture methods except for the high-density spread, which amplified the length bias (**Fig 4b,c**).

**Figure 4.**
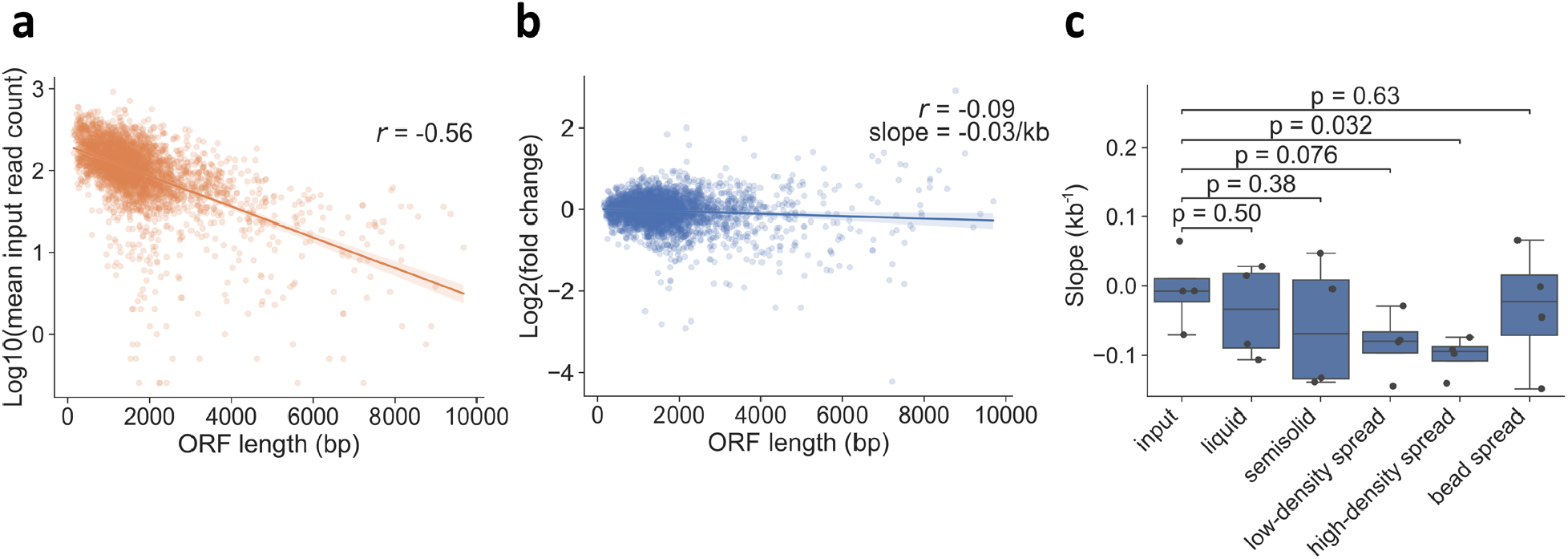
Minimal impact of library culture method on length bias for high coverage transformations. **a**. Comparison of log_10_(mean input read count) (*y* axis) and ORF length (*x* axis) shows the existing length bias in the input MORF library. **b**. Example of how length bias was calculated for each culture condition (for low-density spread replicate 1). The log_2_(fold change) between the sample read counts and average input read counts (*y* axis) was regressed on ORF length (*x* axis) to calculate the slope. **c**. Comparison of abundance-ORF length slopes (*y* axis) for each culture method (*x* axis). Box plot boxes represent the median and interquartile range (IQR). Whiskers extend to points within 1.5 IQRs of the first and third quartiles. P-values were calculated using Welch’s t-test. *r* is the Pearson correlation.

## Discussion

Though plasmid library amplification on agar plates is often used to ensure uniform amplification, we found that plates were at best no better than the simpler liquid culture method for achieving uniform library amplification, and much worse at a colony density that would be reasonable for plating extremely high complexity (>10^8^) libraries. Spreading cells with beads did not improve amplification uniformity in this high colony density scenario. Although amplification in semisolid medium appeared at first glance to produce a less disperse distribution of sequence counts, this method resulted in outlier sequences with counts up to two orders of magnitude above the median count, which was not seen with other culturing methods and could be problematic for certain applications. These overrepresented sequences may have come from the bacteria growing at the top of the culture, which may have had access to more oxygen than colonies deeper in the medium and thus could potentially grow faster, although we tried to mitigate this effect by using tight-sealing lids with little airspace above the media. We also noticed that the yield of cells from semisolid was much lower than from liquid culture (**Fig. S1**). Given these drawbacks, the cost of the specialty agarose required, and the extra effort required to set up the semisolid medium, liquid culture seems to be the best choice for amplifying high complexity plasmid libraries for most applications.

In the case of a lower complexity plasmid library with a wide range of insert sizes, we found that there was almost no measurable difference between culture methods when sufficient library coverage was used (∼100X). The change in barcode count distribution and number of dropped-out barcodes was minimal between the input library and the amplified libraries, and no length-specific bias in plasmid amplification was observed for any culture method except for high-density spread, where, counter intuitively, the bias was amplified. This suggests that changes in library sequence representation during amplification are largely due to random differences in the growth of individual transformant clone populations, which can be averaged out with sufficient coverage during transformation.

Our results suggest that the extra time and cost involved in more complicated semisolid or plate-based culture techniques is unnecessary for many plasmid libraries, as the simplest method, liquid culture, performs at least as well for uniform library amplification. While the plate-based methods looked no better (and often worse) by almost every measure, it is possible that libraries containing sequences causing strong fitness defects (e.g. protein toxins) could still benefit from plate-based culture methods, but investigation of these special cases is beyond the scope of our study.

## Methods

### MORF Input Library Preparation

MORF library without GFP and mCherry controls (Addgene #192821) was transformed into electrocompetent NEB Stable *E. coli* (derived from NEB C3040 and prepared according to Nováková *et al*.^14^) and grown overnight at 30 °C in LB + 100 μg/mL ampicillin. Plasmids were extracted using the GeneJET Plasmid Miniprep Kit (Thermo K0502). Approximately 1.4×10^8^ transformants were obtained, as estimated by colony counting. Library quality was assessed by agarose gel electrophoresis and whole-plasmid Oxford Nanopore sequencing.

### N80 Plasmid Library Construction

An oligo with an 80 nt random region flanked with homology arms for Gibson assembly was ordered from IDT as an Ultramer oligo (**N80_template**). The oligo was double stranded with primer **N80_rev** for a single cycle using Phusion polymerase (NEB M0530). The double stranded N80 library was cloned into the XhoI site of yGPRA_pTpA using Gibson assembly and purified with SPRI beads. At this point each N80 sequence in the library had an equal abundance of one.

### Library Amplification

For each library, all samples were prepared from a single pool of electroporated cells to ensure a similar number of transformants across replicates. Electrocompetent cells were prepared according to Nováková *et al*.^14^ for both experiments. NEB Stable *E. coli* was used for the MORF library, while *E. coli* DH10B was used for the N80 library. Growth temperature was 30 °C for the MORF library and 37 °C for the N80 library.

All culture media contained 100 μg/mL ampicillin and were prewarmed to the growth temperature. Plates were prepared with either 20 mL (90 mm diameter plates) or 56 mL (140 mm diameter plates) of LB agar. Liquid cultures were grown in 10 mL LB in a 50 mL conical plastic centrifuge tube. Semisolid cultures were grown in 15 mL of medium in a 15 mL conical plastic centrifuge tube. Semisolid medium was prepared by adding 0.3% w/v ultra-low gelling temperature agarose (Sigma A5030) to LB and autoclaving. Before use, the medium was boiled to melt the agarose, then cooled to the growth temperature.

After transformation of each library, SOC medium was added to a total volume of 5 mL. Cells were recovered at the required growth temperature for 1 hour with shaking, then placed on ice to prevent further growth. From 5 mL of recovered cells, cultures were started as follows:

- 200 μL was transferred to 10 mL of LB.
- 200 μL was transferred to 15 mL of semisolid medium and mixed thoroughly by inverting. The tube was placed on ice for 15 minutes to solidify the medium.
- 200 μL was spread on a 90 mm plate using a glass spreader until all liquid was absorbed.
- 200 μL was spread on a 90 mm plate with 10 ColiRollers Plating Beads until all liquid was absorbed.
- 66.7 μL was diluted in 489 μL SOC, then plated on a 140 mm plate using a glass spreader until all liquid was absorbed. This was repeated two more times for a total of 200 μL of recovered cells plated across three plates.

This was repeated three more times from the same sample of recovered cells for a total of four replicates. Plates and semisolid medium were incubated in a static incubator. Liquid culture was incubated with shaking at 250 rpm. MORF cultures were grown for 26 hours, and N80 cultures for 16 hours. Serial dilutions were plated to estimate the number of transformants per sample. The estimated number of transformants per sample was ∼3.7×10^5^ for the MORF experiment and ∼7.6×10^4^ for the N80 experiment.

Cells were harvested from plates by scraping three times with LB, ensuring that all colonies were harvested. The harvested cells were vortexed vigorously to ensure that colonies were broken apart and cells were mixed evenly. Liquid and semisolid media were harvested by centrifugation at 4000 × g.

Plasmids were purified from harvested cells using a standard alkaline lysis miniprep procedure and quality was verified by agarose gel electrophoresis. Plasmid concentrations were normalized according to concentrations measured by absorbance at 260 nm prior to sequencing library preparation.

### Sequencing

Sequencing libraries were generated by PCR with NEBNext Ultra II Q5 Master Mix (NEB M0544) in two consecutive reactions. qPCR was used to determine the optimal number of cycles to use. The barcode regions of each plasmid library were first flanked with Illumina TruSeq read 1 and read 2 sequences by PCR using **MORF/N80_TruSeq_i5** and **MORF/N80_TruSeq_i7** primers. Libraries were then indexed using custom ordered TruSeq indexing primers (**Index_i5 and Index_i7**). Indexed libraries were purified by freeze-and-squeeze gel extraction followed by SPRI bead purification. Pooled indexed libraries were sequenced on an Illumina NovaSeq X Plus with 150 bp paired-end reads.

**Table 1.**
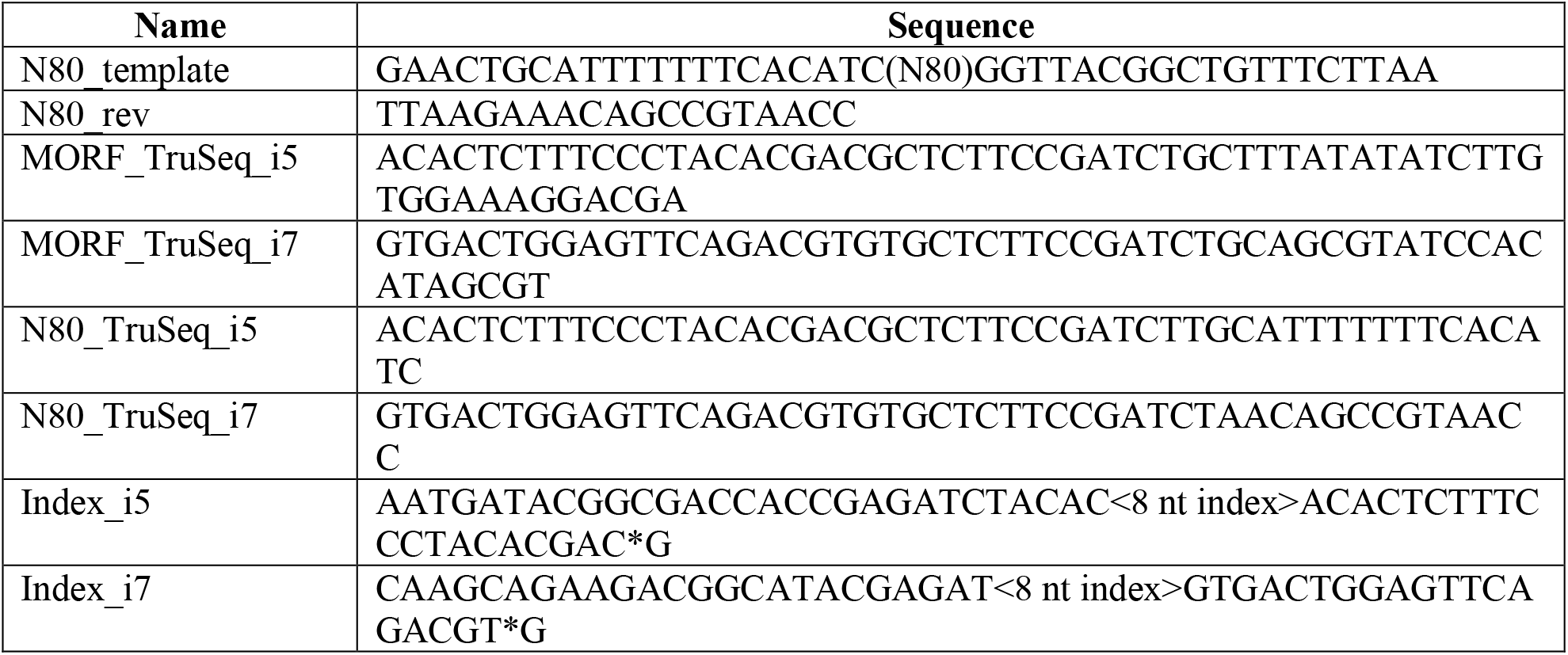
Oligos used. ‘*’ indicates a phosphorothioate bond.

### Processing

The MORF and N80 sequencing data were processed similarly. Each sample was randomly sampled to the lowest read count observed across all samples within each experiment using seqtk (version 1.3) (https://github.com/lh3/seqtk). The barcode sequences were extracted from reads using Cutadapt (version 4.1)^15^ in paired end mode, discarding any reads that were not the expected size (24 bp for MORF and 80 bp for N80) after trimming. Read pairs were merged using NGmerge (version 0.3)^16^. To account for PCR and sequencing errors, merged reads were clustered with starcode (version 1.4)^17^ using default Levenshtein distance of 3 and the connected components clustering algorithm (-c option). We considered a cluster centroid to be a true starting sequence present in the library before transformation, and the number of cluster components the read count for that sequence. The Snakemake^18^ pipeline used for all processing steps can be found at https://github.com/de-Boer-Lab/PlasmidLibraryCulture.

### Analysis

Jupyter notebooks containing all analysis code can be found at https://github.com/de-Boer-Lab/PlasmidLibraryCulture. Pearson correlation between sample and input counts for the MORF library was calculated as follows: for each replicate, a Pearson correlation was calculated between each sample’s counts and the mean of the input counts for the other three replicates. This was done to make input versus input correlations comparable to the others. To determine length bias, for each replicate, first the log_2_(fold change) between the read counts of each sample and the mean read counts of the input library for the other three replicates were calculated. The log_2_(fold change) was then regressed on ORF length (kb) using ordinary least squares linear regression to obtain a slope value that represents the degree of length bias.

## Supporting information

yGPRA_pTpA plasmid

## Data Availability Statement

The data underlying this study are openly available in the SRA at BioProject accession PRJNA1116148. Processed read count data are available at https://github.com/de-Boer-Lab/PlasmidLibraryCulture.

## Supporting Information

Comparison of *E. coli* pellet yield from each culture condition and images of each library culture method (PDF).

Sequence of the yGPRA_pTpA plasmid used for N80 library construction in GenBank format (TXT).

## Author Information

### Authors

**Nicholas Mateyko** - Genome Science and Technology Graduate Program, University of British Columbia, 2222 Health Sciences Mall, Vancouver, BC, Canada.

## Acknowledgments

This research was supported by the Natural Sciences and Engineering Research Council of Canada (RGPIN-2020-05425), the Stem Cell Network (ECR-C4R1-7), and the Canadian Institute for Health Research (PJT-180537, PJT-185988). This research was enabled in part by support provided by WestGrid (westgrid.ca) and the Digital Research Alliance of Canada (alliancecan.ca), and Advanced Research Computing at the University of British Columbia. N.M. was supported by NSERC CGS M and UBC 4YF. The authors wish to acknowledge Canada’s Michael Smith Genome Sciences Centre, Vancouver, Canada for sequencing services. Multiplexed Overexpression of Regulatory Factors (MORF) Library was a gift from Feng Zhang (Addgene #192821).

