## Supporting Information for "Culture wars: Empirically determining the best approach for plasmid library amplification"

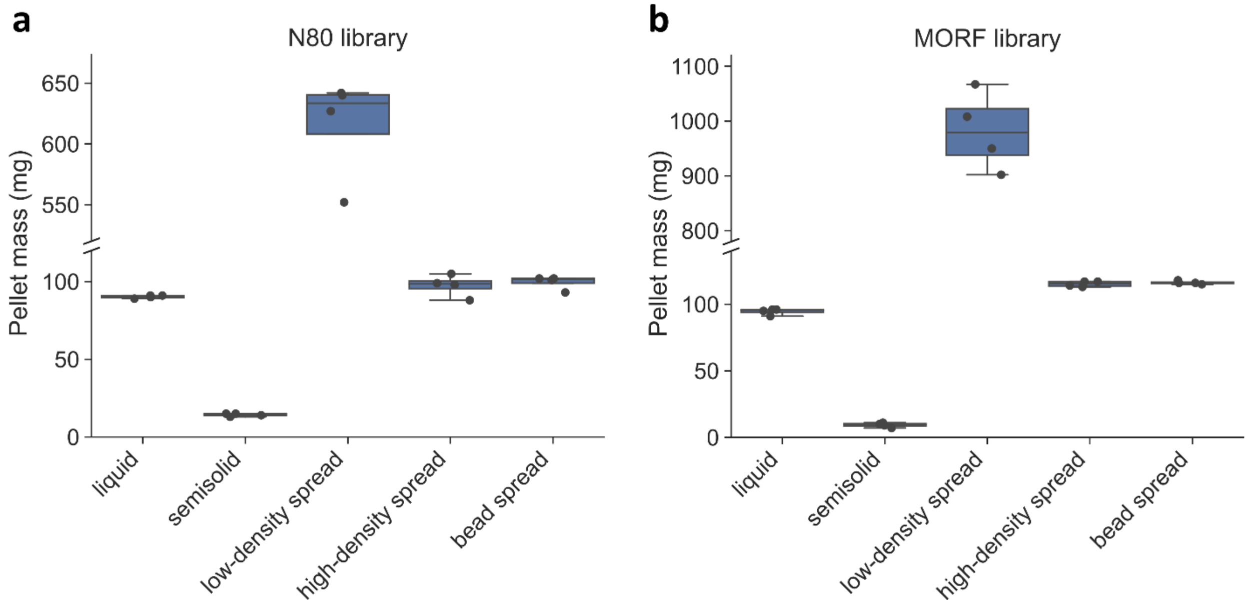


**Figure S1. Semisolid culture produces low cell yield.** Total *E. coli* cell pellet mass (*y* axis) from each culture method (*x* axis) for the (**a**) N80 library and (**b**) MORF library. The *y* axis is broken to better visualize the difference between low-density spread (which used more media per transformant) and other culture methods.

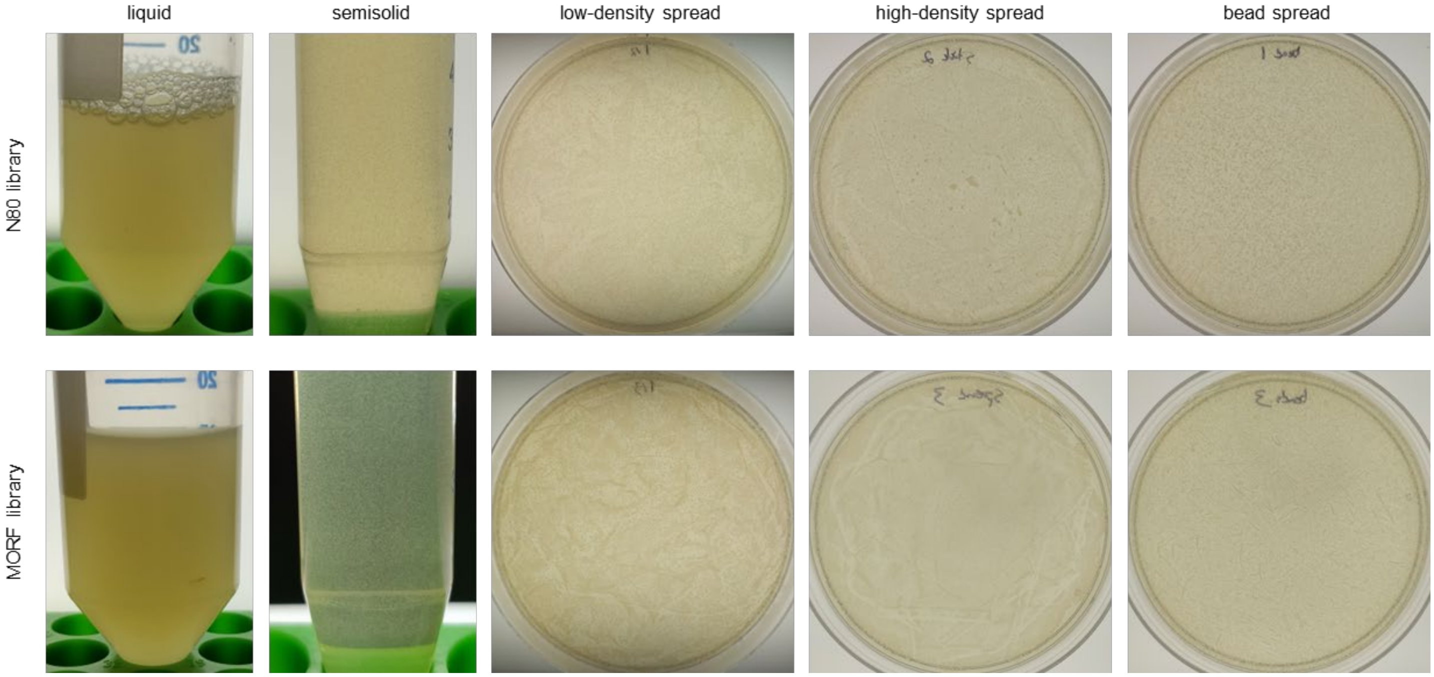


**Figure S2. Library culture images.** Representative images of each library culture method for the N80 and MORF libraries. Low density spread plate is 14 cm diameter, while high-density and bead spread were 9 cm diameter.
